# Differential gene expression in populations of sympathetic neurons purified from a neuropeptide Y reporter mouse

**DOI:** 10.1101/128074

**Authors:** Kristine M. Sikora, Paul H. M. Kullmann, John P. Horn

## Abstract

Using gene chip methodology, we identified candidate genes differentially expressed by vasomotor and non-vasomotor sympathetic neurons in the superior cervical ganglion. Groups of 10 neurons were manually sorted after isolation from a transgenic neuropeptide Y (NPY) reporter mouse that expresses the humanized *Renilla* green fluorescent protein (GFP) under control of Npy promoter sequences. Anatomical analysis of GFP and NPY co-expression showed that 98% of GFP-positive neurons and 16% of GFP-negative neurons express NPY. The probability of contamination in sorted cell samples was therefore high. To minimize this problem, we screened amplified cDNA samples by the quantitative polymerase chain reaction. This approach identified 101 candidate genes preferentially expressed by GFP-positive neurons and 74 candidate genes preferentially expressed by GFP-negative neurons.

## 1. Introduction

Functional subsets of sympathetic neurons in paravertebral chain ganglia are defined by the peripheral target tissues that they selectively innervate, by their central connections and by the differential expression of neuropeptides and other markers (Gibbins, 1995; Janig et al., 1992). For example, distinct groups of sympathetic neurons innervate blood vessels, the heart, glands, pilomotor hairs and other targets, while neuropeptide Y (NPY) is preferentially expressed by vasomotor and cardiac neurons and by smaller subsets of neurons that innervate the pineal gland and iris (Corr et al., 1990; Gibbins, 1991; Gibbins, 1992; Horn et al., 1987; Li et al., 2006; Lundberg et al., 1984; Lundberg et al., 1983; Reuss et al., 1989). The utility of NPY as a neuronal marker in the sympathetic system arises from the high level of protein expression that permits relatively easy detection in cell bodies and peripheral axons by numerous commercially available antibodies. Despite these advantages, NPY expression does not distinguish subsets of sympathetic neurons that innervate different vascular beds. For example, it would be useful to have markers that identify neurons that selectively regulate blood vessels in the brain, skin, heart, kidneys and striated muscle. Such markers could serve as tools for experimentally targeting functional components of the sympathetic motor system to investigate their normal roles in blood pressure regulation and thermoregulation. They could also help to identify pathophysiological changes in animal models of hypertension, diabetes and Parkinson’s disease. As a first step towards that goal, the present study sought to discover new candidates for genetic markers that distinguish NPY-positive from NPY-negative neurons.

Our approach employed a transgenic reporter mouse that expresses the humanized *Renilla* variant of green fluorescent protein (GFP) together with sequences that promote expression of the Npy gene (van den Pol et al., 2009). The strategy was to dissociate ganglia, manually sort neurons into fluorescent and non-fluorescent groups and then assay them using Illumina bead-chip technology. A key reason for using NPY as the marker for cell sorting was that 50-70% of the neurons in paravertebral ganglia at all segmental levels express NPY (Garcia-Arraras et al., 1992; Gibbins, 1995; Headley et al., 2005; Horn et al., 1987; Lundberg et al., 1982). In addition, the structure of NPY is highly conserved from lower vertebrates up through birds and mammals (Larhammar, 1996). This marker thus provides a practical tool for dividing sympathetic neurons into 2 large groups with ancient evolutionary lineages tied to the regulation of cardiovascular and non-cardiovascular targets. The present analysis focused upon the superior cervical ganglion (SCG) at the rostral end of the paravertebral chain because it has been studied more extensively than other chain ganglia and because it is the largest ganglion with the most cells. Even so, the 20,000 neurons found in the SCG do not contain enough RNA for direct detection with gene chip methodology. Instead, we employed an RNA amplification strategy starting with small groups of 10 neurons (Okaty et al., 2011a; Okaty et al., 2011b). We validated the reporter construct by analyzing the co-expression of NPY and GFP in tissue sections and then checked the efficacy of sorting by testing samples for NPY mRNA expression using quantitative real-time PCR (qPCR) prior to gene chip analysis.

## 2. Materials and methods

All animal protocols for this project were approved by the Institutional Animal Care and Use Committee at the University of Pittsburgh. Transgenic Npy-GFP (B6.FVB-Tg(Npy-hrGFP)1Lowl/J) male mice and C57BL/6 wildtype females were purchased (#006417, Jackson Laboratory, Bar Harbor, ME) and then bred by crossing heterozygote with wildtype individuals. Dr. Bradford B. Lowell originally generated the Npy-GFP line (van den Pol et al., 2009). These mice express a bacterial artificial chromosome containing humanized *Renilla* green fluorescent protein under control of upstream and downstream sequences that contain the Npy promoter. Mice were genotyped from tail biopsies taken between postnatal days 14 and 16. Genomic DNA was extracted with the Gentra Puregene Mouse Tail Kit (#158267, Qiagen, Valencia, CA) according to the manufacturer’s protocol. PCR amplification of target regions from 20 ng of genomic DNA used the following reaction mix: 5Uμl Taq DNA polymerase (#18038042 Invitrogen, Grand Island, NY), 10 mM dNTP mix (#10297018, Invitrogen) and 10 μΜ primers (Integrated DNA Technologies, Inc., Coralville, IA). Thermocycling parameters and primer sequences followed genotyping protocols in The Jackson Laboratory database.

The primer sequences were:

Common 5′-TATGTGGACGGGGCAGAAGATCCAGG-3′

Wild type Reverse 5′-CCCAGCTCAC ATATTTATCTAGAG-3′

Mutant Reverse 5′-GGTGCGGTTGCCGTACTGGA-3′

PCR products were separated by gel electrophoresis on 1.5% sodium-borate agarose gels containing 0.5 μg/ml ethidium bromide and visualized with a UV transilluminator.

### 2.1 NPY immunocytochemistry and quantification

SCGs were dissected from six, 3-5 month old heterozygous Npy-GFP mice (males and females), followed by overnight immersion at 4°C in Zamboni’s fixative containing 2% paraformaldehyde and 0.2% picric acid in 0.2 M phosphate buffer. Fixed SCGs were washed in phosphate buffered saline (PBS) and incubated overnight at 4°C in 30% sucrose in PBS. Cryoprotected tissue was embedded in OCT compound (Tissue-Tek 4583, Sakura, Torrance, CA) and cut into 10 μm cryostat sections. Non-adjacent sections were collected from each SCG, blocked and permeabilized by incubation for 30 minutes at room temperature in PBS with 10% normal donkey serum and 0.5% Triton X-100 (PBSDT). Sections were then incubated overnight at 4°C in PBSDT with primary NPY antibody (either 1:3,000 rabbit monoclonal α-NPY (#11976S, Cell Signaling Technology, Danvers, MA) or 1:1,000 sheep polyclonal α-NPY (#AB1583, EMD Millipore, Billerica, MA). After washing with PBS, sections were incubated for 2 hours at room temperature in secondary antibody diluted in PBSDT (either 1:200 Cy3 AffiniPure donkey α-rabbit IgG (#711-165-152, Jackson ImmunoResearch Laboratories, West Grove, PA) or 1:200 Cy3 AffiniPure donkey α-sheep IgG (#713-166-147, Jackson ImmunoResearch Laboratories)). The sections were then washed with PBS, coverslipped and imaged using a Zeiss Axioskop2 microscope, AxioCam HRc camera and AxioVision software. Although superior tissue histology and NPY immunoreactivity can be obtained by embedding tissue in polyethylene glycol and cutting at room temperature (Headley et al., 2005), this method proved unsatisfactory because it destroyed GFP fluorescence.

### 2.2 Neuronal dissociation and sorting

SCGs dissected from four month old (microarray 1) and seven month old (microarray 2) heterozygous Npy-GFP male mice were placed in chilled L-15 media (#SH30525.01, Thermo Scientific). Under a dissection microscope, the ganglia were desheathed, cut into pieces and incubated for 30 minutes at 37°C in pre-warmed L-15 containing 2 mg/ml collagenase type 4 (#LS004186, Worthington Biochemical, Lakewood, NJ), followed by incubation for 30 minutes at 37°C in pre-warmed L-15 containing 0.25% trypsin (#15050-057, GIBCO, Grand Island, NY), with gentle agitation every 10 minutes. To stop the digestion, trypsin was neutralized by diluting 1:10 with growth media that contained MEM (#SH30601.01, Thermo Scientific), 10% fetal bovine serum (#S11150, Atlanta Biologicals, Norcross, GA), 1% penicillin-streptomycin (#B21210, Atlanta Biologicals), 10 ng/mL nerve growth factor (# BT-5017, Harlan Bioproducts, Indianapolis, IN), 0.4 μM cytosine arabinoside hydrochloride (#C6645, Sigma-Aldrich). After pelleting the tissue by mild centrifugation for 1 minute at 100 x g, excess media was aspirated, and the ganglia were triturated in 300 μl of media using three flame-polished Pasteur pipettes with openings of decreasing diameter. We then plated the cells on two 35mm glass-bottom dishes fabricated with 14 mm collagen-coated coverslips (#P35GCOL-1.0-14-C Mattek Corp, Ashland, MA), allowed them to adhere to the substrate by incubating them at 37°C, 5% CO_2_ for 1 hour and began the process of manual sorting.

Sorting was done by placing each 35 mm dish onto a Zeiss IM 35 inverted microscope and identifying GFP positive and negative neurons under epifluorescence illumination. Individual adherent phase-bright neurons were drawn into patch style glass pipettes with 30-60 μ openings using gentle suction and transferred into tubes containing 2 μl of direct lysis buffer (from the Ovation® One-Direct System, see below). The tubes were flash-frozen in liquid nitrogen and stored at -80°C before processing. Collecting each sample of 10-12 cells took approximately 30 minutes. Transfer pipettes were fabricated from 100 μl disposable borosilicate pipettes with a P- 87 puller (Sutter Instruments, Novato, CA), mounted in a suction holder designed for patch-clamp electrophysiology and controlled with a Leitz mechanical micromanipulator.

### 2.3 RNA Isolation, cDNA synthesis and amplification

Groups of cells stored in direct lysis buffer were thawed on ice, followed by cDNA synthesis and amplification using the Ovation® One-Direct System (#3500-12, NuGen, San Carlos, CA) according to the manufacturer’s protocol.

### 2.4 qPCR For Npy

Npy expression was quantified in amplified cDNA samples using the SYBR® Green detection system (#4309155, Life Technologies). 20 μl qRT-PCR reactions included 250 nM primers and 5 ng of cDNA template. Each reaction was run in triplicate on an Applied Biosystems Prism 7900HT cycler with the default real-time PCR program, followed by melt curve analysis.

Primers sequences were:

Mouse Gapdh Forward 5′-CATGGCCTTCCGTGTTCCTA-3′ (IDT, Custom)

Mouse Gapdh Reverse 5′-CCTGCTTCACCACCTTCTTGAT-3′ (IDT, Custom)

Mouse Npy Forward 5′-CCAGACAGAGATATGGCAAGAG-3′ (IDT, Custom)

Mouse Npy Reverse 5′-GGGTCTTCAAGCCTTGTTCT-3′ (IDT, Custom)

### 2.5 Labeling of amplified cDNA and gene chip analysis

Amplified single-stranded cDNA products were purified using the MinElute Reaction Cleanup Kit (#28204, Qiagen, Germantown, MD) according to the Ovation® One-Direct System protocol and stored at -20°C. The University of Pittsburgh Genomics Research Core (www.genetics.pitt.edu) performed quality checks on cDNA samples, labeled them and ran the gene chip assays. cDNA quality was assessed using the Agilent Bioanalyzer and an RNA 6000 Nano chip. cDNA was labeled using the Encore BiotinIL Module (#4210-48, NuGen). Gene expression assays used the MouseWG-6 v2.0 Expression BeadChip (#BD-201-0202, Illumina, San Diego, CA).

### 2.6 Data analysis software

Data analysis utilized Prism 7 (Graphpad, LaJolla, CA), Igor 6 (Wavemetrics, Lake Oswego, OR) and Microsoft Excel. Microarray data analysis employed BRB-ArrayTools (http://brb.nci.nih.gov/BRB-ArrayTools/), developed by Dr. Richard Simon and BRB-ArrayTools Development Team.

## 3. Results

### 3.1 Overlap between NPY and GFP expression

The original work first describing the NPY reporter mouse showed good correspondence between neuronal NPY and GFP expression in the hypothalamic arcuate nucleus (van den Pol et al., 2009), but did not examine sympathetic ganglia. Before studying differential gene expression in sympathetic neurons sorted by GFP expression, it was therefore important to assess co-expression of the reporter with NPY. Figure 1 illustrates the co-localization of NPY-immunofluorescence and GFP in sections of the SCG. As expected, most sympathetic neurons that expressed NPY (Fig. 1A) also expressed GFP (Fig. 1B) and most NPY-negative neurons were GFP-negative. However, the match between expression of the neuropeptide and reporter was not perfect. Figure 1C illustrates an NPY-negative cell that was GFP-positive (Fig. 1D) and Fig. 1E illustrates a NPY-positive cell that was GFP-negative (Fig. 1F). In order to quantitate these observations, fluorescent and non-fluorescent cells were photographed and counted in 7 ganglia from 4 mice. Twenty-one microscope fields were analyzed in non-adjacent sections to avoid double counting of split cells. In 2,423 cells, 53% (1,282 neurons) were NPY-positive and 47% (1,141 neurons) were NPY-negative, which confirms previous work (Gibbins, 1991). However, only 45% of neurons (1099/2423) expressed the GFP reporter, while 55% were GFP-negative (1324/2423). To understand the disparity between NPY and GFP expression, we examined the incidence of double labeling. All but 22 of the GFP-positive cells were NPY-positive. Hence there was a 0.98 (1077/1099) probability that a GFP-positive cell correctly reported the expression of NPY. This contrasted with 205 NPY-positive neurons that failed to express the GFP reporter. The probability of a GFP-negative cell also being NPY-negative was therefore only 0.84 (1117/1322). In other words, GFP expression reliably reported NPY expression 98% of the time, but the absence of GFP was less reliable in that it only reported the absence of NPY 84% of the time. This failure to observe GFP in 16% of NPY-positive neurons could arise either from a GFP detection problem, or from incomplete penetrance of the transgenic GFP reporter construct, possibly due to novel regulatory elements for the NPY gene that are absent in the GFP transgene construct. In an effort to distinguish these possibilities, we measured the distributions of cell body size in the different cell groups, which can reflect functional specialization of sympathetic cell types (Dodd et al., 1983; Gibbins, 1991; Li et al., 2006).

**Figure 1.**
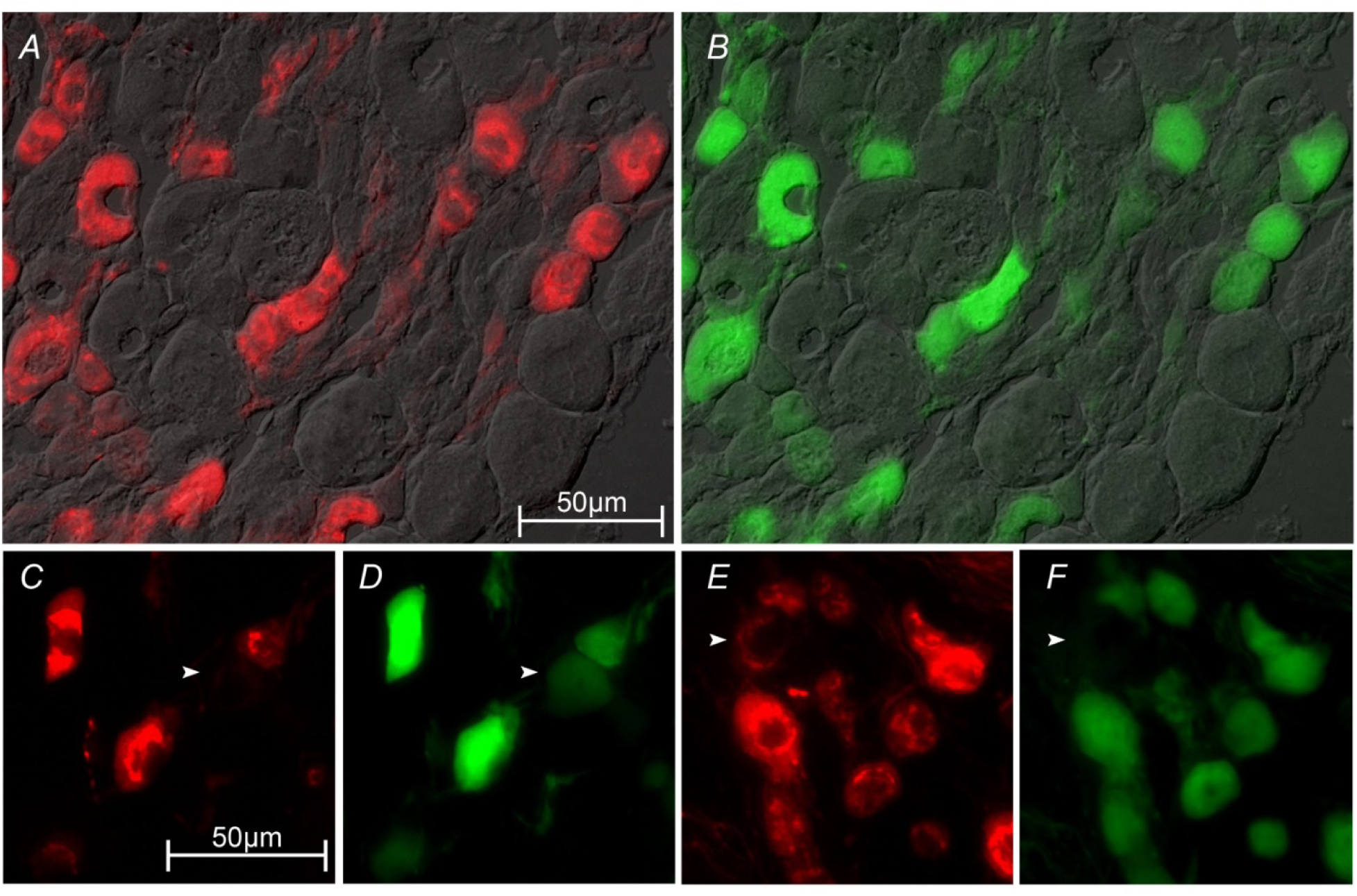
Comparison of NPY-immunoreactivity (*A*,*C*,*E*) with GFP expression (*B*,*D*,*F*) in the superior cervical ganglion. The images in *A* and *B* illustrate the strong correspondence between expression of NPY and GFP in sympathetic neurons from the transgenic mouse model. Unstained cells in the background were photographed using Nomarski differential interference contrast illumination. These panels also show that the largest sympathetic neurons tend not to express either marker. White arrowheads in *C* and *D* point to a rare example of a sympathetic neuron that was NPY-negative and GFP-positive. The arrowheads in *E* and *F* point to a NPY-positive neuron that was GFP-negative.

As shown in Figure 1A, NPY-positive sympathetic neurons tended to be smaller than NPY-negative neurons. In the rat SCG, this is due in part to large secretomotor neurons that project through the external carotid nerve to salivary glands (Headley et al., 2005). To compare the sizes of the different cell groups, we traced and compared the areas of the 2,423 cell bodies whose GFP fluorescence and NPY-immunoreactivity had been scored. The double negative neurons were larger on average than cells in the other three groups (Fig. 2A). This was because the largest cells were found only in the double negative group. Comparing the size distributions for double negative and double positive neurons (Fig. 2B) showed they were skewed and had different shapes. We therefore used a non-parametric ANOVA that detected differences in the 6 possible pairwise comparisons between the 4 groups (Fig. 2). Post hoc tests revealed statistically significant differences between the double negative and double positive neurons (P<0.0001) and between the double negative and NPY-only neurons (P<0.0001). Plotting all 4 cell size data sets as cumulative probability density functions (Fig. 2C) showed that the three groups expressing either one or both markers were very similar and were distinct from the size distribution of the double negative neurons.

**Figure 2.**
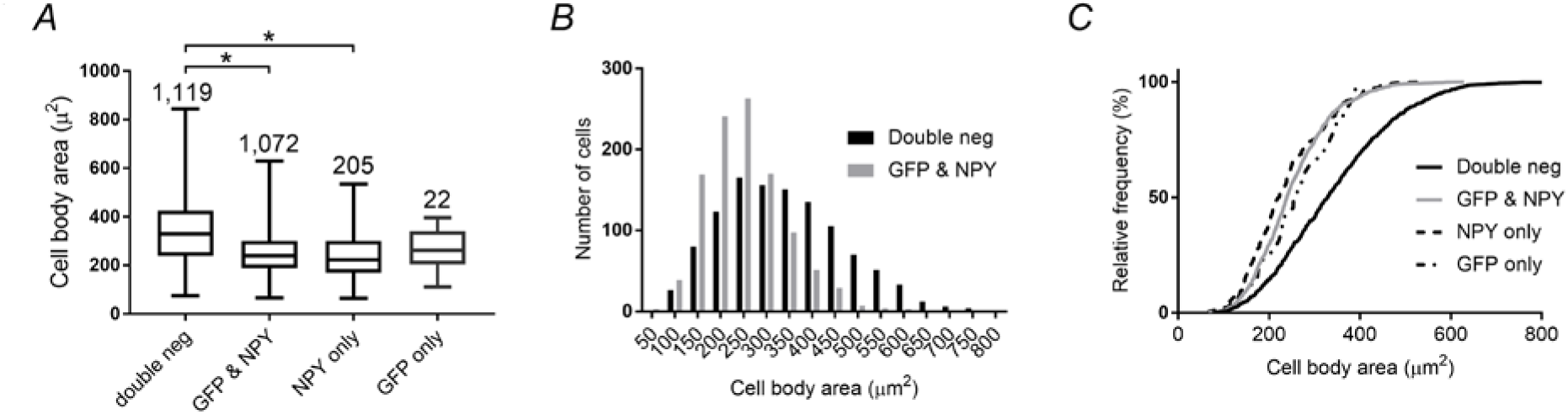
Size distributions of sympathetic neurons that express NPY-immunoreactivity and GFP and of neurons that do not express these markers. Box and whisker plots (*A*) compare the mean, range and 25-75%-tiles of cell body areas for the four groups of neurons. The number of neurons in each group is noted at the top. All groups were compared for statistical differences (* P<0.0001) using a Kruskal-Wallis one-way analysis of variance, followed by Dunn’s multiple comparison test. Histograms (*B*) of cell areas for double positive *and* double negative neurons show that the largest cells are double negative. Normalized cumulative histograms of cell area show that the singly labeled neurons were similar to the double positive cells.

### 3.2 Efficacy of cell sorting

Our experimental design utilized groups of 10-12 neurons in order to increase the signal to noise ratio by smoothing out small variations in gene expression between individual cells (Okaty et al., 2011a). In preliminary gene chip experiments, it was therefore surprising to find that Npy was not always preferentially expressed in groups of GFP-positive neurons. Simple binomial calculations (http://www.graphpad.com/quickcalcs/probability1/) using our staining data helped to explain this result. Given a probability of 0.84 that a GFP-negative cell is also NPY-negative, the probability that 10 GFP-negative neurons picked at random will all be NPY-negative is only 0.174 and the probability that a sample of 10 cells will be contaminated by 1 or 2 NPY-positive neurons is 0.60. Although the odds of avoiding contamination are much better (i.e. 98%) when using GFP to collect NPY-positive neurons, the chance that 10 of 10 cells will be NPY-positive is only 0.817 and there is a 16% chance the sample will have one NPY-negative neuron. As a practical solution to this limitation, we assayed the 10 cell samples used to assess differential gene expression (next section) by qPCR for Npy.

The average cycle thresholds (C_T_) for NPY and GAPDH in 5 samples of GFP-positive cells were 17.26 ± 1.29 and 20.06 ± 0.63 (mean ± standard error). The corresponding (C_T_) values in 5 GFP-negative samples were 22.82 ± 2.39 and 20.56 ± 0.60. Using these numbers to determine relative expression (2^−Δ*C_T_*^ × 1000) showed significantly higher NPY expression in the groups of GFP-positive neurons than in the groups of GFP-negative neurons (Fig. 3). Using the double Δ method (2^−ΔΔ*C_T_*^) (Livak et al., 2001; Pfaffl, 2001), we calculated a 33.3 fold difference in NPY expression levels between the two groups of sorted neuronal samples.

**Figure 3.**
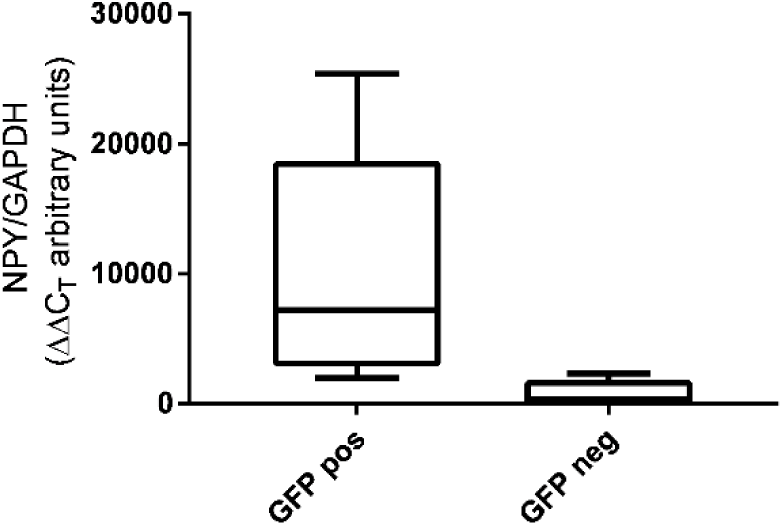
Differential expression of NPY in groups of sympathetic neurons sorted by GFP fluorescence. Amplified cDNA preparations were assayed by qPCR for NPY and GAPDH. The level of NPY expression relative to GAPDH was calculated for each sample from the ΔC_T_. Higher levels of NPY were seen in GFP-positive neuronal groups when compared to the GFP-negative groups (P=0.015, n=5 Mann-Whitney test). Box and whiskers denote the median, range and 25-75%-tiles for each group.

### 3.2 Identification of differentially expressed candidate genes

Twelve neuronal samples from 6 mice were subjected to microarray analysis on two Illumina Beadchip Arrays. Half the samples were GFP-positive and the others GFP-negative. Figure 4 illustrates the raw hybridization signals (range = 0 – 26,651) for each sample, after sorting them in rank order. Data from 2 samples (red) were discarded because the signals were systematically lower than in all the other 10 samples. In the remaining 10 samples, we observed interleaving of signal intensities for the GFP-positive (green) and GFP-negative (blue) data sets, with no systematic bias towards one cell type or the other. Seventy-five percent of the 45,281 probes yielded signals in the range of 80 to 100, with a background signal of about 70. Thus, the majority of probes detected very small responses.

**Figure 4.**
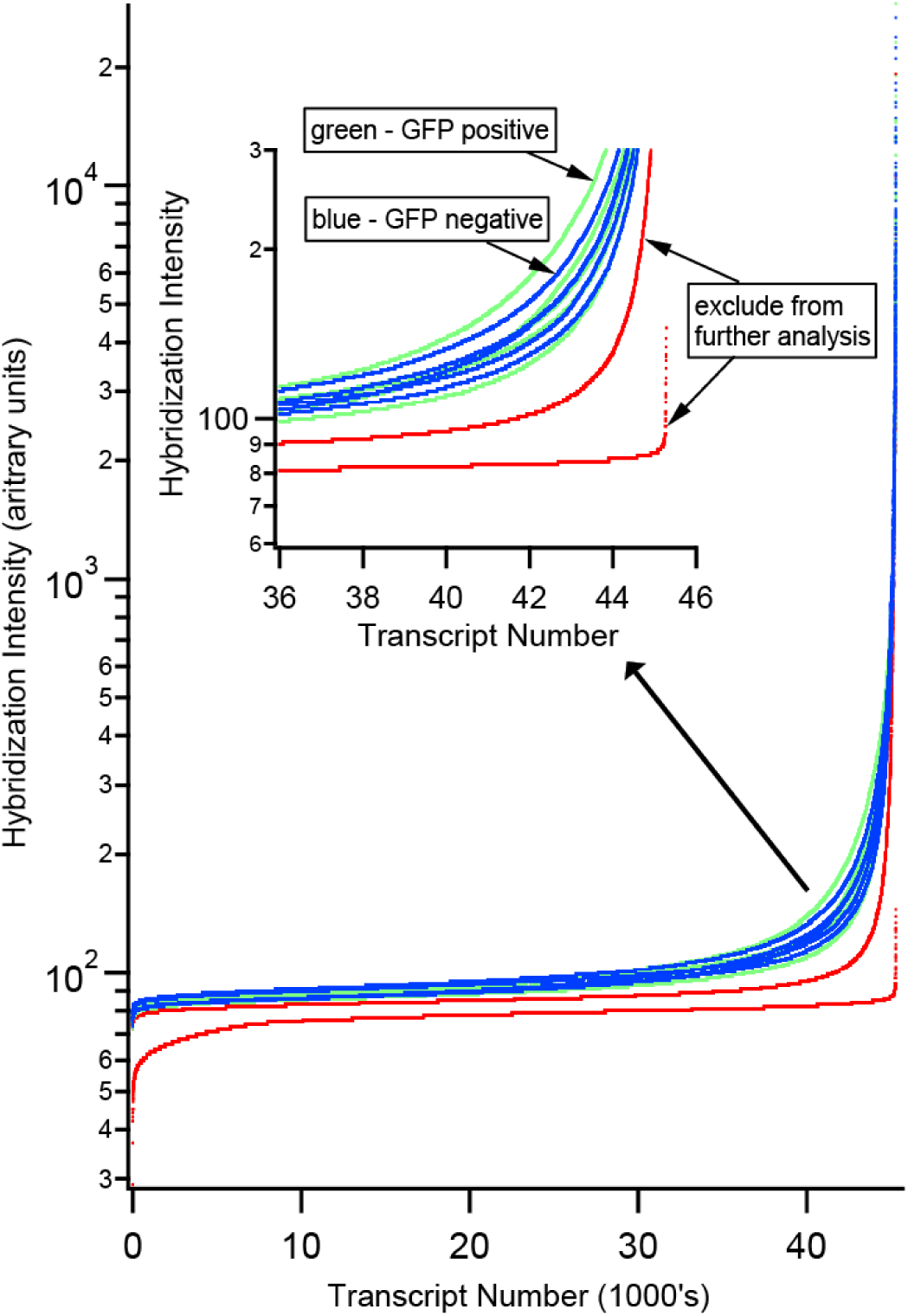
Comparison of hybridization signals from 12 samples run on Illumina bead-chip arrays. Data from each array was plotted in rank order of signal intensity. Two of the samples (red) yielded data that was systematically lower than the others and was removed from further analysis. The other ten samples of GFP-positive (green) cells and GFP-negative (blue) cells, which yielded comparable signal intensities, were analyzed for differential gene expression.

Microarray data from the 10 samples that met our quality control standard of the raw data were imported into BRB-ArrayTools with quantile normalization. Differences in gene expression were determined by comparing normalized gene expression values between GFP-positive samples (JH1, 3, 5, 14, and 22) and GFP-negative samples (JH2, 6, 13, 15, and 23). All of the original and analyzed bead chip data are available on the NCBI/GEO database (Accession GSE81075). With 5 biological replicates in each group, a two-sample t-test was calculated for each gene. A list of 220 genes with P <0.05 was identified as significantly different between the two groups. We then excluded 36 unknown or uncharacterized genes from the list. This left 184 known genes: 104 expressed at higher levels in GFP-positive neurons (Table 1) and 80 expressed at higher levels in GFP-negative neurons (Table 2). Npy at the top of Table 1 had the largest differential expression signal of all genes. This indicates that our strategy for sorting neurons followed by qPCR screening succeeded in comparing pools of cells that were enriched with NPY-positive cells with pools that were largely NPY-negative. Further inspection of the data revealed 2 genes (Akap9, Cdip1) that each came up twice in Table 1, and 5 genes (Enah, Fam110b, Fam3c, Rbm5, Sdhc) that came up 11 times in Table 2. Multiple hits occur because the bead chip methodology includes 45,281 probes that oversample the genome to more complete insure coverage. Taking the duplicates and Npy into account, our data identified preferential detection of 101 new genes in GFP-positive neurons and 74 new genes in GFP-negative neurons.

**Table 1.**
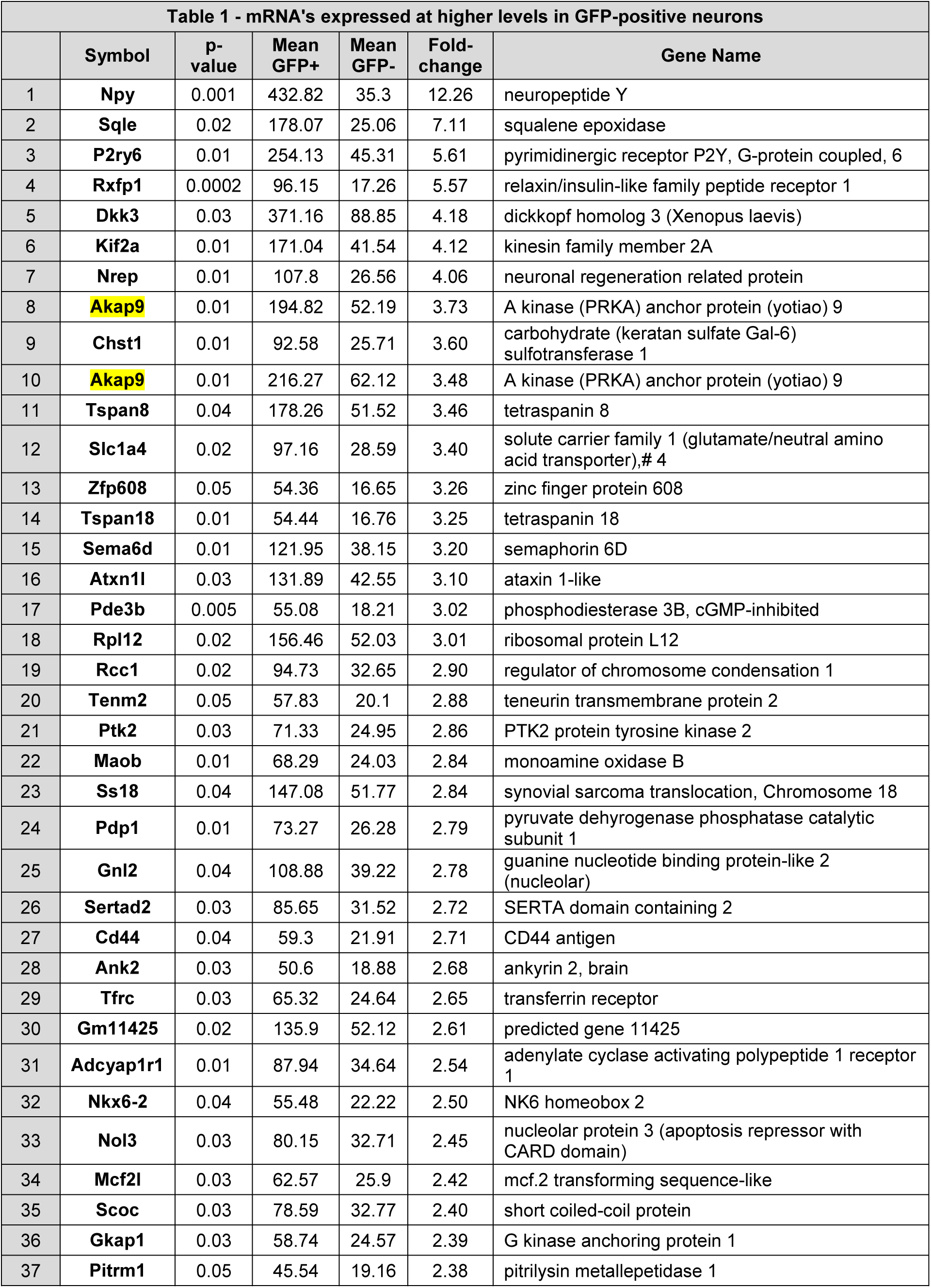
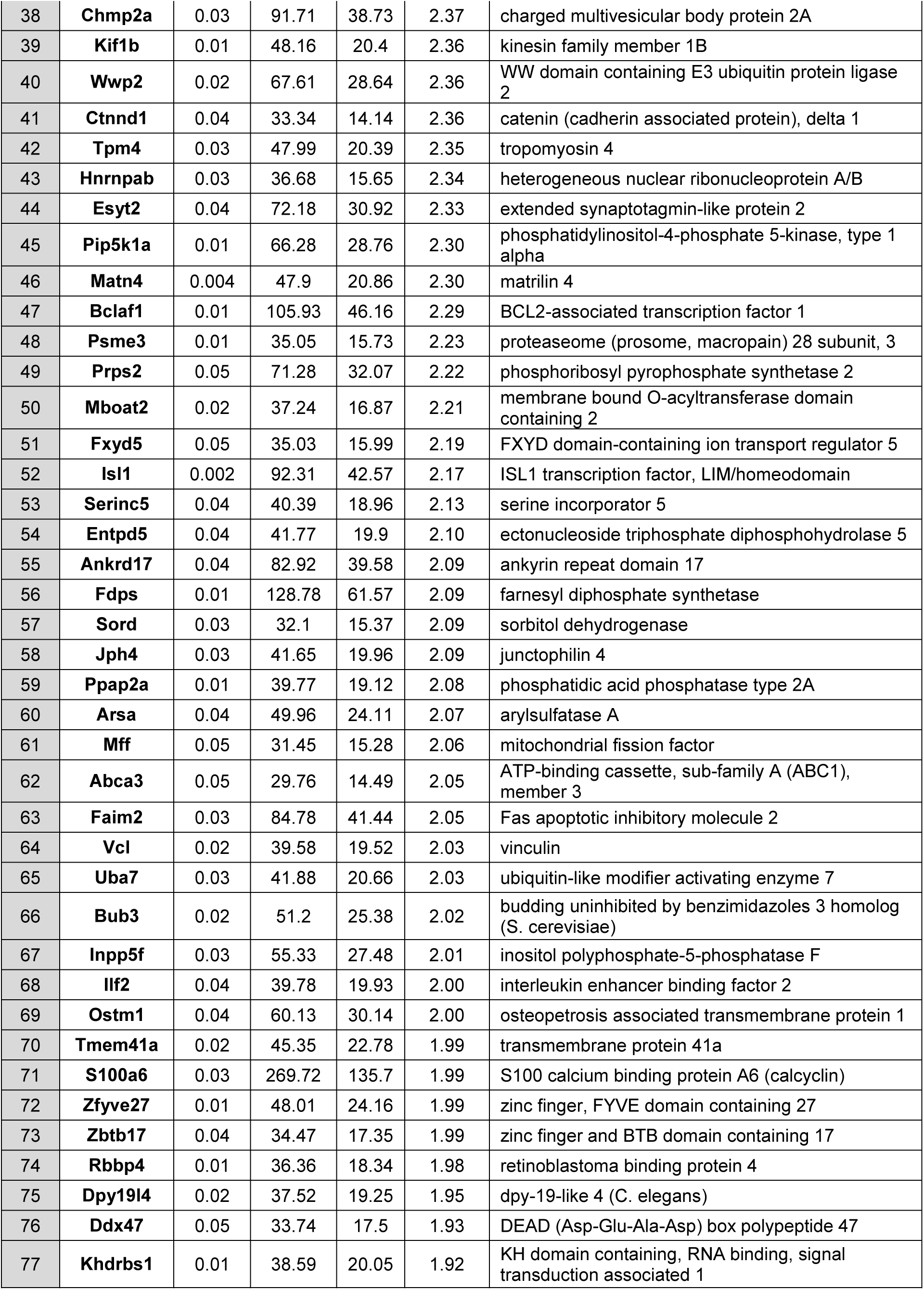
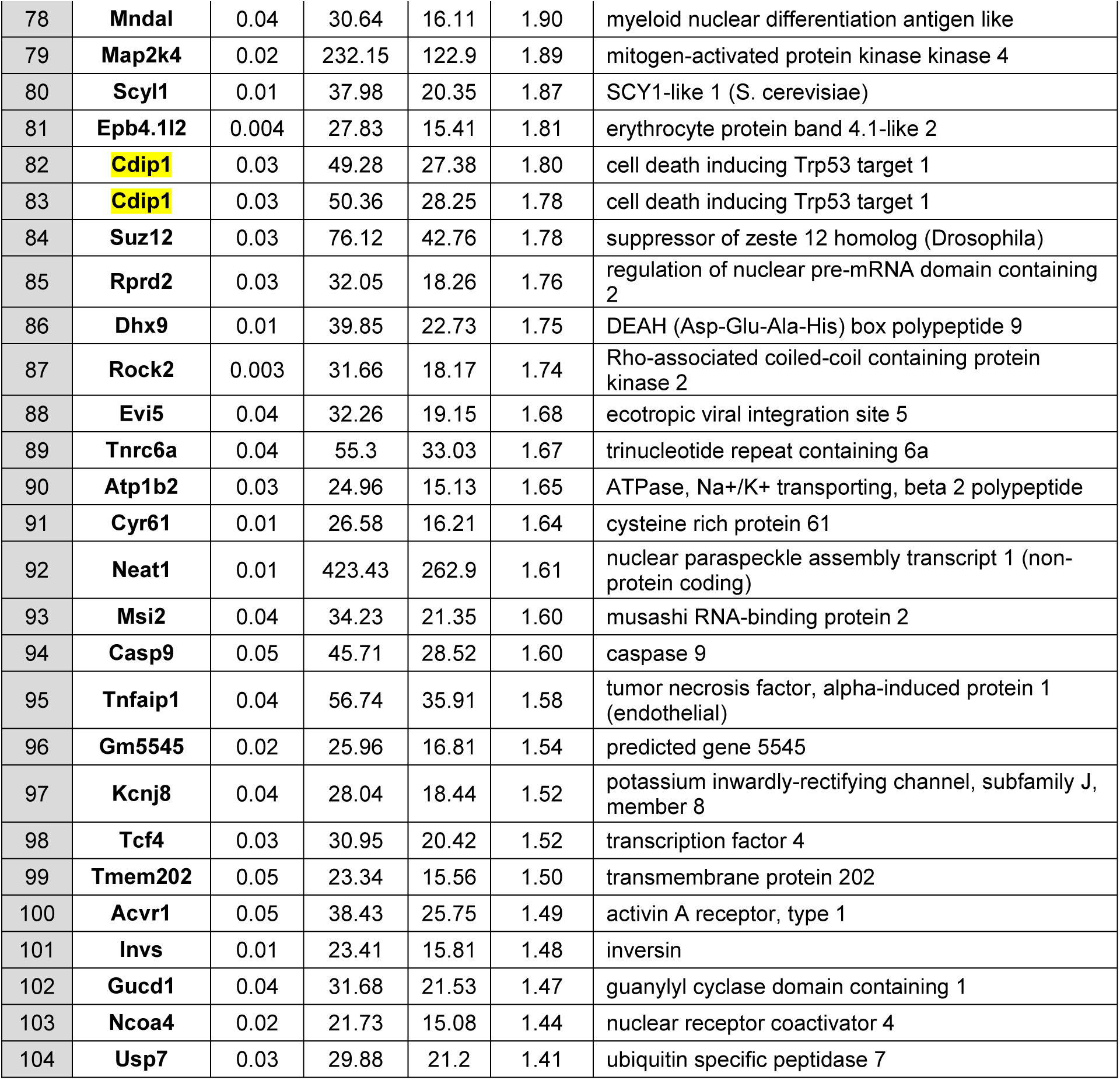
Candidate genes for preferential expression by GFP-positive sympathetic neurons. Accounting for Npy at the top of the list and two genes that appear in duplicate (yellow) yields 101 unique candidates for preferential expression.

**Table 2.**
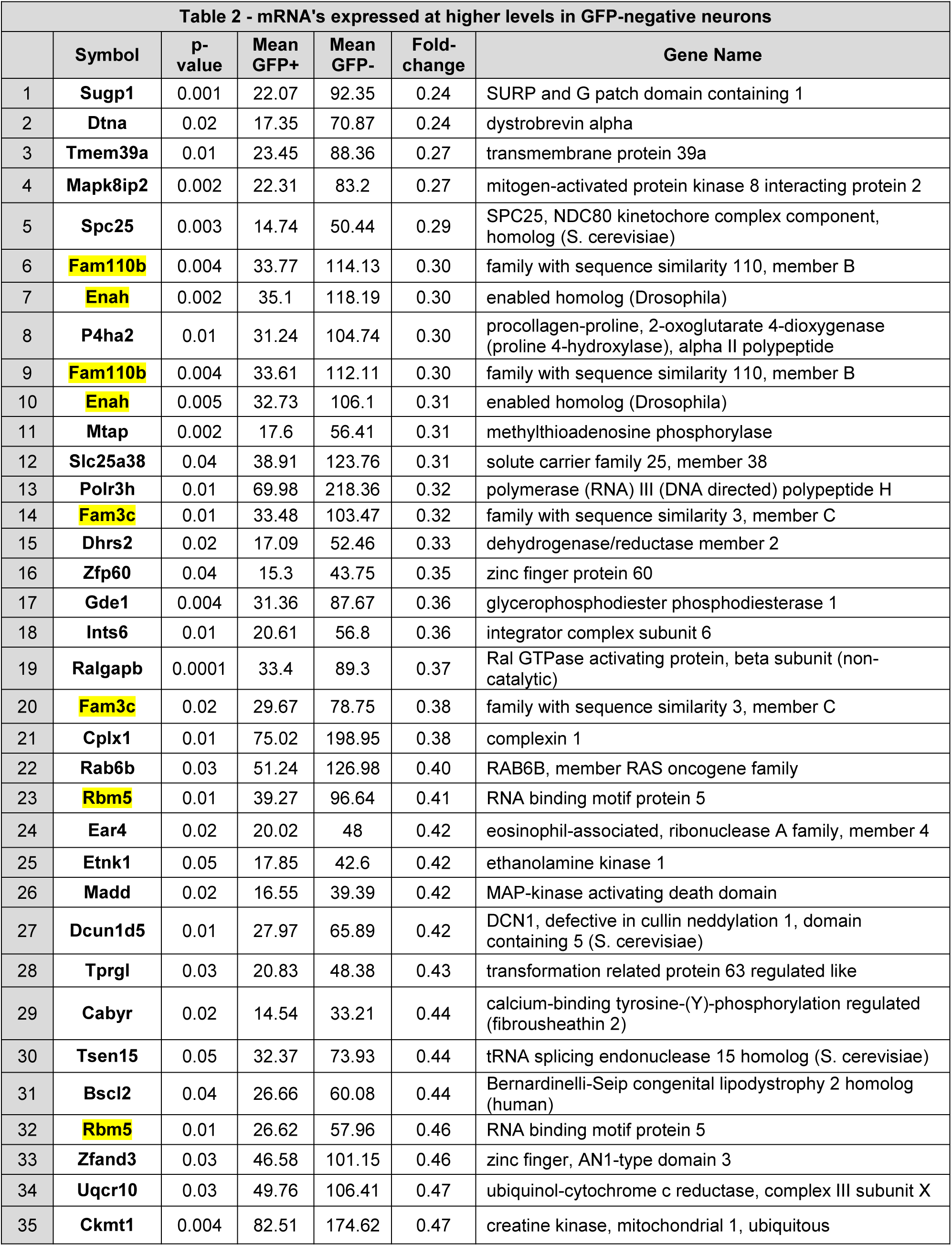
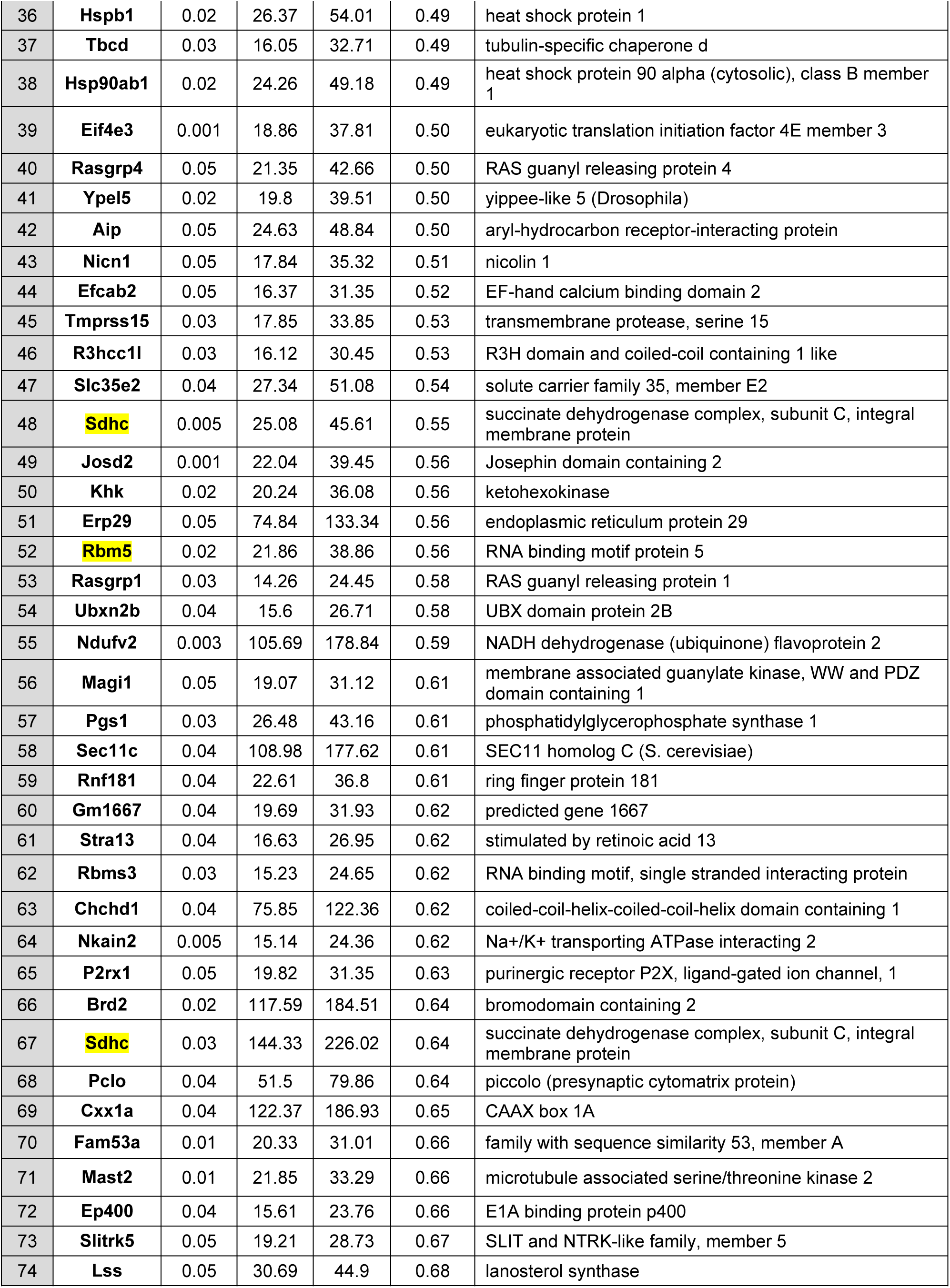
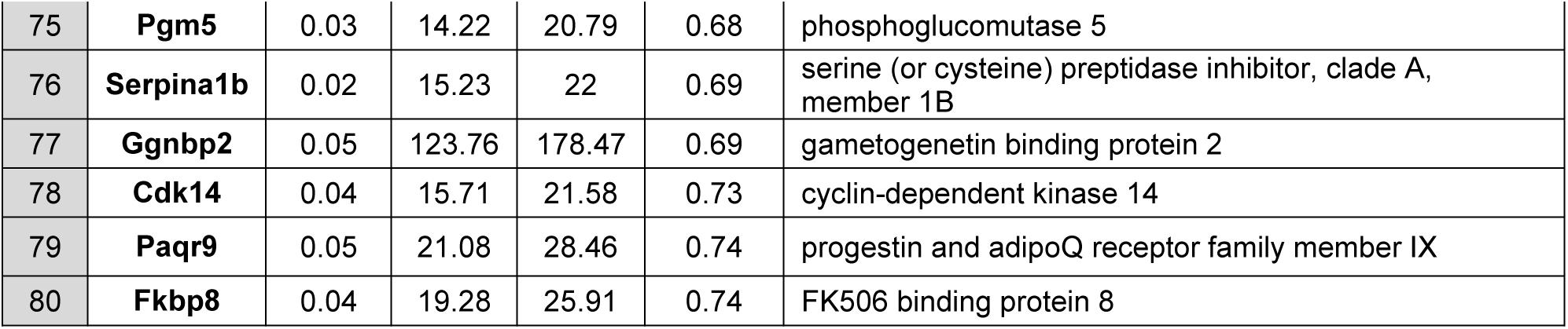
Candidate genes for preferential expression in GFP-negative sympathetic neurons. Five genes are listed multiple times (yellow) resulting in 74 unique candidates.

## 4. Discussion

Reporter mice provide powerful tools for studying specialized types of neurons to understand their physiology and underlying genetics. The Lowell lab developed the Npy-GFP mouse to aid studies of hypothalamic neurons that control feeding and energy homeostasis (van den Pol et al., 2009). Now for the first time, we have used this mouse model to investigate sympathetic neurons, where NPY is a marker for the subset of cells that innervate the cardiovascular system. Comparing the expression of NPY-immunoreactivity and GFP fluorescence confirmed that the transgenic reporter construct used to drive GFP expression is highly effective (Figures 1 and 2). Nonetheless it is not perfect. While 98% of neurons that express NPY also express the GFP reporter, only 84% of neurons that do not express GFP are NPY-negative. This becomes important when sorting neurons to assess differential gene expression. We were able to overcome this difficulty by testing cell samples for differential Npy gene expression using qPCR and by targeting the gene chip analysis to samples that passed this test. The Illumina bead chip array data identified 175 candidate genes that are differentially expressed by NPY-positive and NPY-negative SCG neurons. Many of these genes are familiar as G-protein coupled receptors, as signaling molecules, and as transcription factor related molecules, while others serve in a range of additional capacities. Our efforts to identify molecular networks that might help interpret the candidate’s functional significance have not been successful to this point. It is also important to note that the targets identified remain candidates. Additional studies will be required in order to validate differential expression of these genes. In the meantime, since the completion of this project, the methodology for expression analysis has continued to advance. The bead chip arrays used in our study are now obsolete and no longer available from Illumina. In its place, RNA sequencing methods have become more readily available for the analysis of single cells and large numbers of single cells using microfluidic cell sorting and bar coding methods (Macosko et al., 2015), together with more advanced bioinformatics approaches to the analysis. These and related approaches that employ deep sequencing have now been used to identify neuronal types in the retina (Shekhar et al., 2016), in the hypothalamus (Campbell et al., 2017), in sensory neurons (Chiu et al., 2014) and in sympathetic neurons from thoracic chain ganglia (Furlan et al., 2016). Future work will undoubtedly uncover finer genetic distinctions between subsets of sympathetic neurons that control different facets of autonomic motor behavior and provide tools for analyzing their physiological properties. Although these advances circumvent the need for reporter mice for cell sorting, the Npy-GFP mouse remains promising as a tool for electrophysiological studies of vasomotor sympathetic neurons.

## Acknowledgements

We thank Ms. Deborah J. Hollingshead and Drs. Janette Lamb in the Genomics Research Core Laboratories at the University of Pittsburgh for their advice on experimental design and analysis and for performing the cDNA labeling and gene chip procedures. We also thank our colleagues, Drs. Kathryn Albers, Brian Davis and Rick Koerber, for many helpful discussions.

